# A Reproducible Methodology for High-Resolution 3D and Orthomosaic Mapping in Forested Streams

**DOI:** 10.64898/2026.02.11.705124

**Authors:** Baptiste Nelaton, Eric Harvey, Katrine Turgeon

## Abstract

1 - Frequent, fine-scale assessments of forested stream morphology are essential for understanding habitat conditions and guiding management, yet dense canopy and budget constraints often limit airborne LiDAR and conventional surveys.
2 - We present a reproducible, ground-based photogrammetry workflow using a chest-mounted action camera (GoPro) and the Agisoft Metashape software to generate scaled, georeferenced 3D models and orthomosaics of stream reaches. The workflow details image acquisition under variable lighting and flow conditions, image alignment, dense cloud/mesh generation, and export of GIS-ready products.
3 - Applied in Mont-Tremblant National Park and Kenauk Nature (Québec, Canada) across 56 forest-stream reaches in a single summer field season, the workflow produced scaled, GIS-ready orthomosaics with nominal pixel sizes as low as ∼1.36 mm/pixel. These products captured fine-scale channel morphology, enabling consistent reach-scale habitat mapping and temporal comparisons for restoration monitoring.
4 - By reducing equipment costs and simplifying logistics relative to airborne or terrestrial LiDAR, this practical tool enables frequent, high-resolution surveys in dense-canopy streams. Future extensions (e.g., automated substrate classification via machine learning) can further standardize habitat assessment and support long-term ecological monitoring.

## Introduction

Because of their high sensitivity to perturbations, forest streams are among the first ecosystems to respond to climate change. While water quality and trophic interactions undoubtedly influence the health and productivity of aquatic environments (Patil, 2023; Su et al., 2021; Zhao et al., 2019), the physical dimension of a stream (*e.g.*, bank morphology, bed structure, substrate composition and heterogeneity) is a major determinant of streams’ ability to sustain biological communities and is key to their resilience to disturbances (Heatherly et al., 2007; Mažeika et al., 2004; Rabeni, 2000).

Although stream management and assessment often prioritize biological indicators, these indicators can be highly heterogeneous, context-dependent (*e.g.,* season), and sometimes insensitive to short-term change (Gillmann et al., 2023; Brettschneider et al., 2023; Poppe et al., 2016). Quantifying habitat heterogeneity at the reach scale provides a robust and complementary indicator of habitat quality. Understanding the physical dimension and dynamics of streams across multiple scales can inform management strategies, guide conservation efforts, and evaluate restoration programs.

Forested streams are often hard to access and survey, which increases the need for accurate, lightweight, and adaptable methods to characterize complex and heterogeneous ecosystems. Precisely describing forest-stream habitats therefore requires approaches that produce fine-scale, spatially explicit, and reliable data.

In this context, the acquisition of spatially explicit, high-resolution data has emerged as a key to unravelling the inherent complexity of fluvial systems (Bertin et al., 2015; Hodge et al., 2009). Although approaches to characterize and map river physical attributes exist (Dawson et al., 2024; Dietrich, 2016; Legleiter, 2012; Thoms et al., 2018), they remain underutilized for several reasons. First, they are often described as cumbersome to implement (Dawson et al., 2024; Zhou et al., 2023). Second, they have not kept pace with the increasingly precise surveying tools that have emerged over the past few decades (*e.g.,* drones equipped with high-resolution RGB or multispectral cameras, RTK-GPS systems, LiDAR technologies, and photogrammetric techniques). Finally, the lack of standardized protocols hinders the development of reliable, reproducible diagnostics for both scientific research and ecosystem management.

Photogrammetry, particularly Structure-from-Motion (SfM; estimating 3D structure from a set of 2D images; Eltner & Sofia, 2020), has emerged as a non-invasive, flexible, and affordable solution for acquiring detailed 3D data (Carrivick & Smith, 2019; Leduc et al., 2019; Marteau et al., 2017). SfM leverages overlapping images taken from multiple viewpoints to reconstruct the 3D geometry of a scene without prior knowledge of the camera’s position or orientation. While this approach has proven valuable in various contexts by providing cost-effective high-resolution digital elevation models (DEMs) and 3D models (Westoby et al., 2012), its applications in forested streams remain largely unexplored.

### Purpose of the methodological note

In this methodological note, we present a detailed and replicable protocol for using a chest-mounted GoPro Hero© action camera and Agisoft Metashape photogrammetry software (version 2.0.2) to generate scaled, georeferenced 3D models and orthomosaics of stream reaches.

We first outline the step-by-step procedure for image acquisition to ensure consistent coverage and overlap, even under challenging field conditions. We then describe the image processing workflow in Metashape, from initial photo alignment to the production of a final 3D model and orthomosaic.

To demonstrate the approach in practice, we present a case study using data collected from forested streams in Mont-Tremblant National Park (Québec, Canada), which illustrates the accuracy and ecological relevance of our results (see Supplementary Material for full details). Finally, we discuss the potential applications of these 3D datasets and the advantages and limitations of the proposed method. By offering practical guidance and highlighting key considerations, we aim to facilitate broader adoption of this approach among researchers and environmental managers seeking refined assessments of stream physical structure and habitat complexity.

## Materials and Methods

### 1. Study area and field conditions

The method described is primarily intended for wadeable forested streams, those with dense riparian vegetation, variable channel morphology, and a diverse substrate composition. In practice, this typically includes stream reaches with an average depth of up to 1 m, ensuring safe, manageable conditions for the operator to walk within the channel.

While the approach can be adapted to a range of environmental settings, certain conditions improve data quality and ease of implementation. These include near-baseflow conditions (i.e., low and stable discharge and water level during image acquisition), sufficient ambient light, and favourable weather (e.g., no rain), as well as conditions that minimize turbidity. Although not strictly required, these conditions help produce clearer imagery, improve image alignment, and yield more reliable 3D reconstructions. In suboptimal conditions, such as partially overcast skies or slightly elevated turbidity, data acquisition remains feasible, albeit potentially requiring additional image filtering or processing.

### 2. Equipment used

#### Action Camera

While a variety of action cameras can be used (e.g., DJI Osmo Action 5 Pro©, Insta360 Ace©, Akaso Brave 8©), we used a recent-generation GoPro Hero©. These cameras offer an ultra-wide-angle mode that reduces the number of photographs needed for comprehensive coverage, thereby streamlining data collection. An additional advantage of newer action camera models is the recording of GPS coordinates for each image, which facilitates approximate georeferencing and simplifies subsequent integration of the resulting models into Geographic Information Systems (GIS) platforms.

A pole extension can optionally be used to raise the camera above the water surface, which may improve the field of view and image overlap; however, it is not required for successful reconstructions, and the workflow performs well without it. If the channel and banks lack distinctive topographic features, recognizable objects (e.g., artificial targets or markers printed on paper) can serve as tie points (see Supporting Information, Fig. S1). These easily identifiable markers help ensure accurate image alignment and improve the robustness of the 3D reconstruction.

#### Software

For data processing, we used Agisoft Metashape (v.2.0.2; Agisoft LLC). Several commercial and free alternatives to Agisoft Metashape include RealityCapture (commercial), Pix4Dmapper (drone-focused; commercial), 3DF Zephyr (commercial), and Meshroom (free). These tools (list not exhaustive) offer comparable 3D reconstruction, orthomosaic, and mesh generation capabilities for mapping and modeling but we did not test them. While the Standard Edition of Metashape is sufficient for most SfM workflows, it requires additional steps to produce orthomosaics. The Professional Edition streamlines orthomosaic generation and enhances overall accuracy. Complementary software, such as any GIS platform, can be used in conjunction with Metashape Standard Edition to create orthomosaics and further analyze georeferenced outputs. Processing did not require a high-end workstation; we successfully ran the workflow on a standard laptop, with runtimes depending primarily on image count and dense-cloud quality settings.

### 3. Field data collection protocol

#### Image Acquisition Methodology

The data collection protocol involves walking in the stream (in-channel) along a predefined stream reach and capturing images at regular intervals. Starting from the upstream end, the operator moves downstream, taking a photograph approximately every meter. After reaching the downstream limit, the operator retraces the route back upstream, repeating the photography process at the same intervals. This bidirectional approach increases image overlap and improves the robustness of the 3D reconstruction.

### 3. Data Processing with Metashape

Figure 1 summarizes the six main steps of the data processing approach. The complete and detailed methodology, including specific parameters and settings, is provided in the Supplementary Material (see Supporting Information, Fig. S. 1).

**Figure 1:**
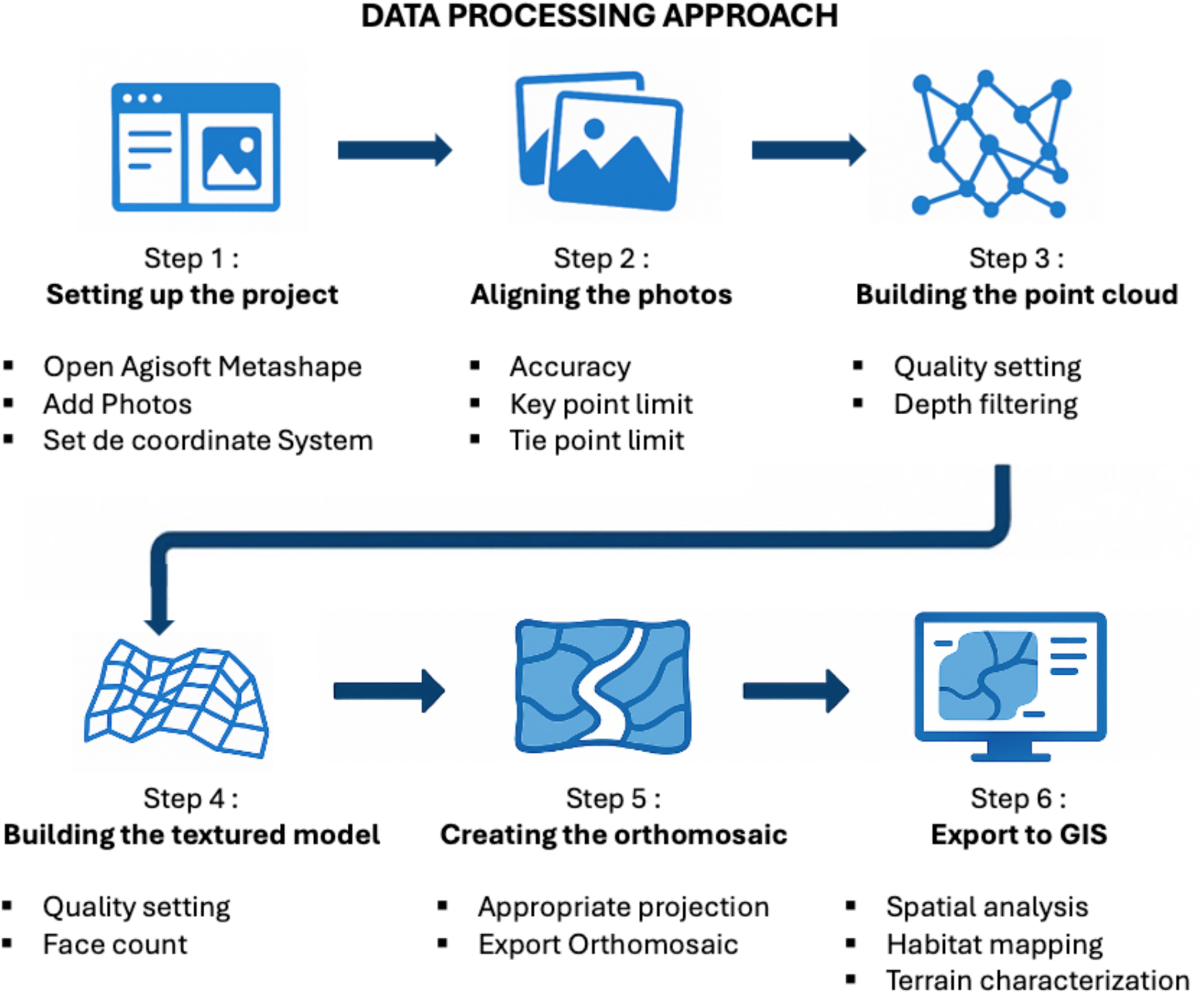
Overview of the photogrammetric workflow used for stream 3D modeling, from image alignment to GIS-based analysis. Detailed processing parameters are provided in Supplementary Material.

## Results

Using ground-level imagery, we produced a high-resolution 3D model and an associated orthomosaic (see Fig. 2 for an example). The final reconstruction achieved a ground sampling distance (GSD) of approximately 1.3 mm/pixel, enabling detailed visualization of channel morphology, substrate composition and heterogeneity, and microhabitat features. Because no independent accuracy assessment was conducted, the reported GSD reflects image resolution rather than absolute positional accuracy. For the Fig. 2 example, field image acquisition required ∼30 minutes, and end-to-end processing on a standard laptop (MacBook, Apple M1) was typically on the order of ∼1 hour for Steps 1–5 (Appendix S2 for full details). We deployed the workflow at scale across 56 wadeable forest-stream reaches during a single summer field season, demonstrating that the protocol can be implemented as a routine monitoring tool.

**Figure 2:**
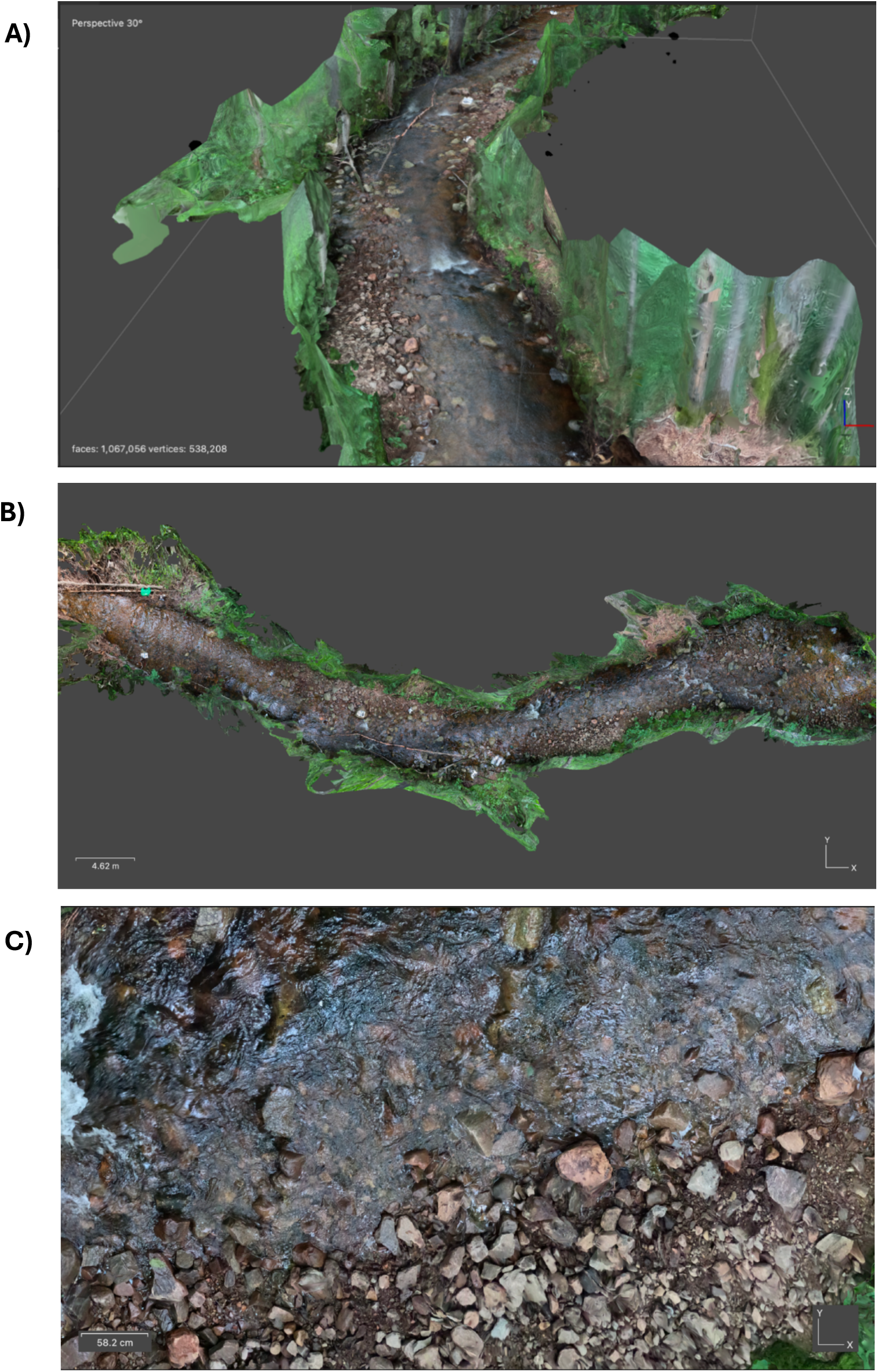
Photogrammetric reconstruction of a forest stream reach using close-range image acquisition. **(a)** Oblique 3D view of the textured mesh model generated in Agisoft Metashape, showing both the channel and streambank vegetation in high detail (1,067,056 faces, 538,208 vertices). **(b)** Top-down orthographic view of the same model, providing a scaled orthomosaic suitable for manual mapping of microhabitats and geomorphic units. **(c)** Close-up view of the orthomosaic at full resolution, illustrating the level of detail available for substrate identification and manual delineation.

## Discussion

### Potential Ecological Interpretations

One of the key strengths of this high-resolution photogrammetry lies in its ability to capture fine-scale features essential to stream ecology. By accurately mapping streambed topography, substrate composition, and physical structures, researchers and managers can derive valuable information from habitat conditions that support aquatic biota at scales relevant to organisms’ perception (e.g., macroinvertebrates to fish). Several key ecological applications include:

#### Habitat characterisation

High-resolution 3-D models generated through photogrammetry provide a richly detailed visual representation of the channel bed, micro-habitats, roughness elements and in-stream refuges (See Supporting Information S.3, Fig. S.2 for an example of stream habitat characterization). Those microhabitats are vital for biodiversity, from diatoms, benthic invertebrates to fish (Nemes-Kókai et al., 2024; Taira et al., 2024; Zhu et al., 2023). At the same time, small-scale roughness elements such as pebbles, cobbles, root wads, and dead wood influence flow patterns and shear stress, shaping micro-habitats heterogeneity across the channel (Wen et al., 2022).

Because this approach is essentially image based it yields only an indirect (i.e., qualitative) quantification of some habitat variables such as water depth, flow velocity and precise substrate type. These variables cannot be extracted automatically from the model itself (at least for now).

#### Quantification of Large Dead Wood (LDW)

Large dead wood significantly influences channel structure, nutrient cycling, and habitat heterogeneity (Cordova et al., 2007). Photogrammetry enables detailed surveys of the size, orientation, and distribution of large dead wood, which can be used to assess its role in forming pools, sorting substrates, and providing cover for aquatic organisms (Andrus et al., 1988; Gurnell et al., 1995).

#### Temporal and Spatial Comparisons

Because photogrammetry datasets are geographically referenced and archived, they can be revisited and compared over time (Gracchi et al., 2021), allowing for longitudinal studies of channel evolution, restoration success, and ecological changes in response to natural events (e.g., floods) or management actions (e.g., LDW enhancement, bank stabilization projects).

#### Integration with Other Ecological and Hydrological Datasets

Once a high-resolution 3D model is generated, it can be combined with other layers of information (Lane et al., 2020) such as fish population data, invertebrate surveys, or riparian vegetation indices to elucidate how physical habitat correlates with biological communities. Additionally, coupling these models with hydraulic simulations (Chen et al., 2019) can identify spatial patterns of flow refugia, spawning habitat, and other ecologically critical areas.

## Methodological Advantages

The presented photogrammetric approach offers several key benefits, making it both accessible and practical for a wide range of end-users, including watershed organizations, governance and national park managers, and academic researchers.

### 1. Ease of Field Implementation

Unlike aerial surveys requiring drones or manned aircraft, this ground-based method relies on minimal equipment and training; a chest-mounted action camera and basic field gear. Its simplicity reduces logistical barriers and ensures that non-specialists can readily adopt the technique. This accessibility expands the potential user base and promotes wider application in both management and research contexts.

### 2. Cost-Effectiveness and Time Savings

Compared to LiDAR campaigns or traditional topographic surveys, the presented approach requires substantially fewer resources. Equipment costs are much lower, and field surveys can be completed more rapidly. By streamlining data collection and processing, practitioners can allocate resources efficiently, enabling more frequent monitoring and evaluation of stream conditions. Moreover, this method provides objective and precise spatial data compared to subjective visual estimations, which are often prone to observer bias and inconsistencies. As a result, the technique supports accurate assessments and reliable decision-making.

### 3. Reproducibility and Long-Term Data Archiving

The method yields spatially explicit, high-resolution datasets that can be easily stored, archived, and revisited over time. This reproducibility facilitates comparative analyses and ensures that high-quality, interpretable visual records are maintained for future reference. As a result, the technique not only supports current management and research objectives but also contributes valuable baseline information for long-term environmental assessments.

## Limitations and Constraints

Despite its clear advantages, the method also faces challenges that may affect data quality and overall usability. First, the stream reach must be safely accessible on foot so that the operator can walk both downstream and upstream while capturing photographs; steep, densely vegetated or hazardous channels may therefore be excluded from analysis. Uneven illumination, strong contrasts, and shadows cast by dense forest canopies can reduce image clarity and complicate point matching, potentially leading to gaps or inconsistencies in the 3D reconstruction. Deep, swift-flowing sections or streambeds with homogeneously colored and textured substrates (e.g., uniform, low-lying green vegetation) pose difficulties for the photogrammetric algorithms. Such conditions may result in fewer discernible features for tie point extraction, diminishing the quality of the model. Overhanging branches and dense riparian vegetation can limit the camera’s line of sight, obstructing views of the streambed. In some cases, minimal clearing may be required to ensure adequate visibility of critical features. While Metashape and modern action cameras offer internal correction routines, residual lens distortions or rolling shutter effects may still occur. Additional calibration steps or the use of carefully selected camera settings can mitigate these distortions, but they remain a potential source of error.

## Conclusion

This methodological note presents a reliable, reproducible protocol for generating scaled and georeferenced 3D models, as well as orthomosaics, of forested stream reaches using ground-based photogrammetry. The resulting high-resolution datasets, combining precise topographic information and advanced modeling techniques, capture the inherent complexity of these environments and thereby enhance our ability to predict and understand hydrological and ecological dynamics. Such detailed spatial information holds significant value not only for aquatic ecology research but also for environmental management, habitat conservation, and long-term restoration efforts. As these methodologies continue to evolve, their integration into long-term ecological monitoring programs promises even greater insights into the temporal variability of stream processes and the effectiveness of management interventions over time.

## Acknowledgements

We thank Parc national du Mont-Tremblant (Sépaq) and Kenauk Nature for site access, logistics, and permits. We are grateful to Gabriel Marcoux-Huard and Mateusz Eugenio-Babinski for insightful field discussions that informed the development of the characterization protocol, and for assistance with pilot surveys and initial processing in Agisoft Metashape. We thank François Degiorgi (Université de Franche-Comté), who introduced us to the IAM habitat characterization method, which helped shape our thinking on the importance of habitat in stream ecology.

**Use of AI tools:**OpenAI ChatGPT (model GPT-5; accessed August 2025) was used only for language editing. All suggestions were reviewed and implemented by the authors; the corresponding author takes responsibility for the final text.

## Funding

This work was supported by the Groupe de Recherche Interuniversitaire en Limnologie (GRIL).

## Author contributions

Baptiste Nelaton, Katrine Turgeon, and Éric Harvey conceived the study. Baptiste Nelaton developed the methodology, conducted fieldwork, performed the analyses, and prepared the visualizations. Katrine Turgeon and Éric Harvey supervised the project and secured resources/funding. Baptiste Nelaton drafted the manuscript; all authors revised it critically and approved the final version.

## Conflict of interest disclosure

The authors declare that they have no financial conflicts of interest in relation to the content of the article. The authors declare no non-financial conflicts of interest.

## Data, scripts, code, and supplementary information availability

A minimal reproducible example dataset (inputs, settings, and outputs, including the Metashape project and processing report) is available on Zenodo: DOI 10.5281/zenodo.18924673.

## SUPPORTING INFORMATION

## Appendix S1. Ground-control targets and placement along the reach (field photo)

**Fig S1:**
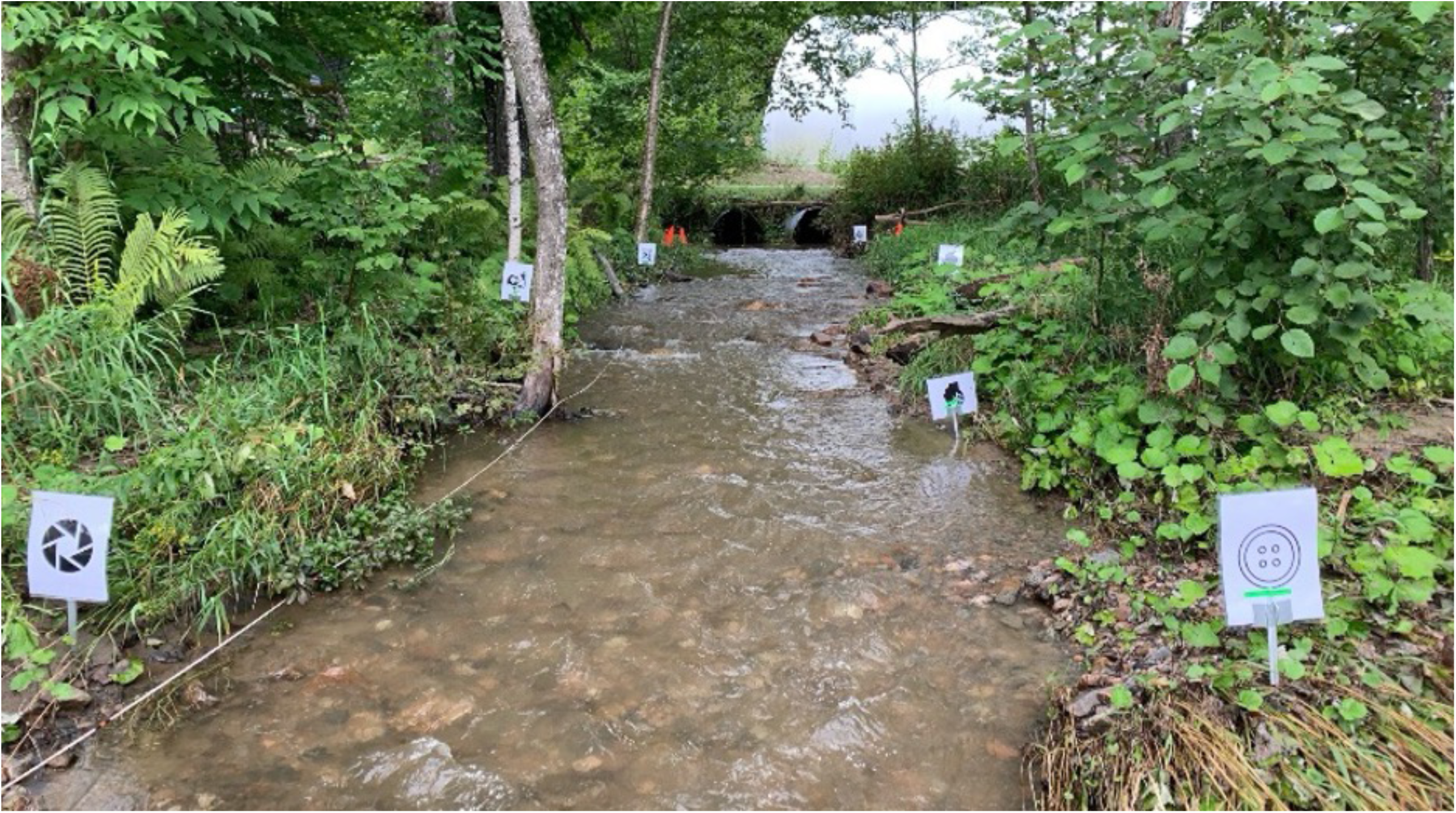
Ground-control targets (GCPs) placed along a stream reach to aid image alignment and georeferencing in close-range photogrammetry.

## Appendix S2. Detailed workflow: a step-by-step guide to ground-based photogrammetry of forested streams (Agisoft Metashape)

**1 - Data Processing with Metashape**

**Step 1: Setting Up the Project (**∼**10min)**

1. **Open Agisoft Metashape :**

◦ Launch the software on your computer.
2. **Add Photos:**

◦ Go to **Workflow > Add Photos**.
◦ Select all the images captured with the action camera.
◦ Click **Open** to import the images into the project.
3. **Set the Coordinate System:**

◦ If your images include georeferencing information or you are working in a known coordinate system, select it under **Reference Settings**.
◦ For data from Quebec, consider using an appropriate projected coordinate system such as **NAD83 / Quebec Lambert** (EPSG:32198) if it suits your study area.

**Step 2: Aligning the Photos** (∼**15 min)**

1. **Photo Alignment Settings:**

◦ Go to Workflow > Align Photos.
◦ Under Accuracy, select High (a good balance between processing time and accuracy; choose Highest if you have a powerful machine and want the best possible alignment).
◦ Key point limit: Set this to around 80,000 to ensure sufficient feature detection.
◦ Tie point limit: Set this to about 4,000 to maintain a manageable number of tie points without overwhelming the project. These settings typically provide a good balance: the key point limit is high enough to capture fine details of the stream bed and banks, while the tie point limit helps control for noise and reduce processing overhead.
2. **Start the Alignment:**

◦ Click **OK** to begin photo alignment.
◦ Wait for the alignment process to complete. Depending on the number of photos and system performance, this step may take approximately 10–20 minutes.

**Step 3: Building the Point Cloud (∼30 min)**

1. **Build Point Cloud:**

◦ Go to **Workflow > Build Point Cloud**.
◦ Choose a **Quality** setting. We recommend **Medium** quality for a reasonable trade-off between processing speed and detail. Higher settings (e.g., High) offer more detail but significantly increase computation time.
◦ **Depth Filtering:** Select **Mild** depth filtering. This setting usually strikes a good balance between smoothing out noise and retaining important details in complex stream environments.
2. **Run the Process:**

◦ Click **OK** to start building the Point cloud.
◦ Allow the process to complete. Processing times vary based on image count and computer performance.
3. **Save the Project:**

◦ Go to **File > Save As**.
◦ Name your project (e.g., “DSM_Model”) and click **Save**.

**Step 4: Building the Textured Model (∼15 min)**

1. **Build Mesh (if desired):**

◦ **Workflow > Build Mesh**
◦ Use the Point Cloud as the source.
◦ Select a **Face Count** (e.g., Medium) to produce a detailed yet manageable mesh.
2. **Build Texture:**

◦ **Workflow > Build Texture**
◦ Use default settings or adjust as necessary for better texture quality.
◦ Click **OK** to generate the textured model.

**Step 5: Creating the Orthomosaic (∼10 min)**

1. **Build Orthomosaic:**

◦ Go to **Workflow > Build Orthomosaic**.
◦ If you have georeferenced images or GCPs, choose the appropriate **Projection** (e.g., NAD83 / Quebec Lambert).
◦ If no georeferencing is available, select a **Planar** projection (Top XY) to generate a relative (non-georeferenced) orthomosaic.
2. **Start the Process:**

◦ Click **OK** to begin.
◦ Wait for the orthomosaic generation to complete.
3. **Complete and Save:**

◦ Once finished, **File > Export Orthomosaic** to your preferred format and location.

Always use a coordinate system appropriate for your region and data sources to ensure better accuracy in analyses. Higher-quality settings (Highest accuracy, High dense cloud) demand more processing power and time. If your hardware is limited, opt for lower-quality settings. If initial results are unsatisfactory, consider adding artificial targets, adjusting camera angles, or using different key/tie point limits to improve alignment and reconstruction quality.

## Appendix S3. Integrating close-range photogrammetry with morphodynamic habitat assessment in a forested stream

Because this approach is essentially image based (the analyst inspects the orthomosaic or shaded-relief model on screen and traces the boundaries by hand) it yields only an indirect (i.e., qualitative) quantification of habitat heterogeneity: depth, flow velocity and substrate type cannot be extracted automatically from the model itself (at least, not for now).

Nevertheless, when photogrammetric mapping is coupled with complementary field protocols, such as the Indice d’Attractivité Morphodynamique (IAM; Degiorgi et al. 2002), which scores habitat quality from the diversity and attractiveness of depth, velocity and substrate classes for fish, the spatial precision of those indicators is greatly improved. The high-resolution polygons produced in the GIS can be overlaid on IAM survey points to refine class boundaries, verify transitions between sequences like run - riffles - pools, and ensure that coarse elements (pebbles, cobbles, root wads, woody debris) that influence shear stress are properly accounted for (see example figure).

In this way, photogrammetry transforms a point-based field index into a spatially explicit map, enhancing our ability to identify reaches that would benefit from added cover or structural enhancement and to design interventions that maintain (or restore) habitat complexity and ecological resilience.

**Figure S.3:**
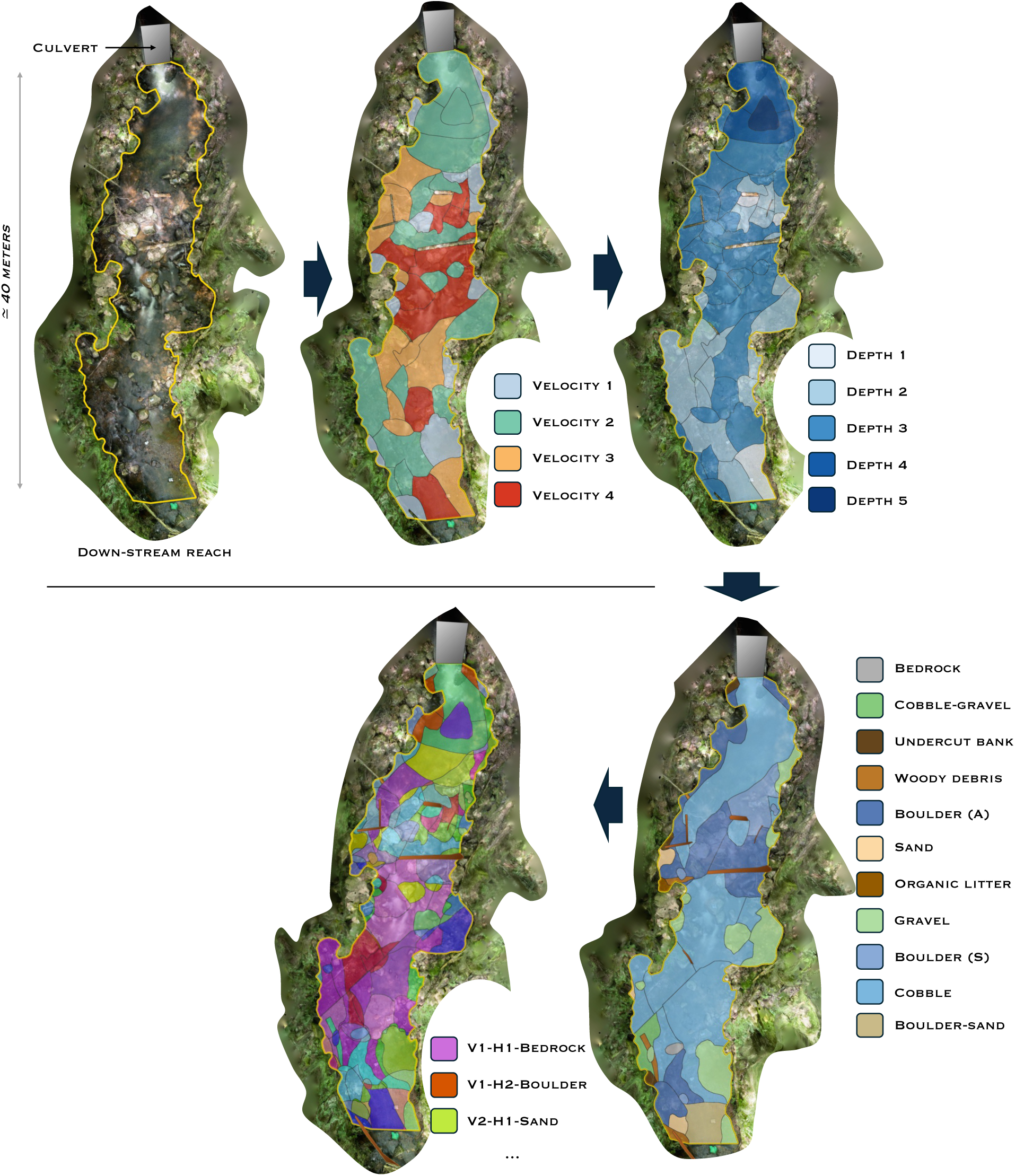
Workflow for combining photogrammetric mapping with morphodynamic habitat assessment (IAM protocol) in a forested stream. Up: Orthomosaic generated from close-range photogrammetry, providing a high-resolution aerial view of the stream reach. Middle: Delineation of channel contours and boundaries in a Geographic Information System (GIS) environment based on the orthomosaic. Down: Integration of photogrammetric data and field-based IAM (Indice d’Attractivité Morphodynamique) protocol to map fish habitat attraction zones. Each polygon represents a spatial unit classified by combinations of depth, flow velocity, and substrate characteristics relevant to fish habitat suitability.

## Notes

### Competing Interest Statement

The authors have declared no competing interest.

### Summary of Updates

Added a supplementary information availability section, with a Zenodo deposit providing a minimal reproducible example dataset (DOI 10.5281/zenodo.18924673).

## Bibliography

1. Agisoft LLC (2024) Metashape Professional, version 2.0. Agisoft LLC, St. Petersburg, Russia. Available at: https://www.agisoftmetashape.com

2. Andrus, C. W., Long, B. A., & Froehlich, H. A. (1988). Woody Debris and Its Contribution to Pool Formation in a Coastal Stream 50 Years after Logging. Canadian Journal of Fisheries and Aquatic Sciences, 45(12), 2080–2086. 10.1139/f88-242

3. Bertin, S., Friedrich, H., Delmas, P., Chan, E., & Gimel’farb, G. (2015). Digital stereo photogrammetry for grain-scale monitoring of fluvial surfaces: Error evaluation and workflow optimisation. ISPRS Journal of Photogrammetry and Remote Sensing, 101, 193–208. 10.1016/j.isprsjprs.2014.12.019

4. Carrivick, J. L., & Smith, M. W. (2019). Fluvial and aquatic applications of Structure from Motion photogrammetry and unmanned aerial vehicle/drone technology. WIREs Water, 6(1), e1328. 10.1002/wat2.1328

5. Chen, Y., DiBiase, R. A., McCarroll, N., & Liu, X. (2019). Quantifying flow resistance in mountain streams using computational fluid dynamics modeling over structure-from-motion photogrammetry-derived microtopography. Earth Surface Processes and Landforms, 44(10), 1973–1987. 10.1002/esp.4624

6. Cordova, J. M., Rosi-Marshall, E. J., Yamamuro, A. M., & Lamberti, G. A. (2007). Quantity, controls and functions of large woody debris in Midwestern USA streams. River Research and Applications, 23(1), 21–33. 10.1002/rra.963

7. Dawson, C., Ashmore, P., & Corry, R. (2024). Evaluating river morphology change with a geomorphic form variation approach. Earth Surface Processes and Landforms, 49(2), 855–874. 10.1002/esp.5744

8. Dietrich, J. T. (2016). Riverscape mapping with helicopter-based Structure-from-Motion photogrammetry. Geomorphology, 252, 144–157. 10.1016/j.geomorph.2015.05.008

9. Eltner, A., & Sofia, G. (2020). Structure from motion photogrammetric technique. In Developments in Earth Surface Processes (Vol. 23, pp. 1–24). Elsevier. 10.1016/B978-0-444-64177-9.00001-1

10. Gracchi, T., Rossi, G., Tacconi Stefanelli, C., Tanteri, L., Pozzani, R., & Moretti, S. (2021). Tracking the Evolution of Riverbed Morphology on the Basis of UAV Photogrammetry. Remote Sensing, 13(4), 829. 10.3390/rs13040829

11. Gurnell, A. M., Gregory, K. J., & Petts, G. E. (1995). The role of coarse woody debris in forest aquatic habitats: Implications for management. Aquatic Conservation: Marine and Freshwater Ecosystems, 5(2), 143–166. 10.1002/aqc.3270050206

12. Heatherly, T., Whiles, M. R., Royer, T. V., & David, M. B. (2007). Relationships between Water Quality, Habitat Quality, and Macroinvertebrate Assemblages in Illinois Streams. Journal of Environmental Quality, 36(6), 1653–1660. 10.2134/jeq2006.0521

13. Hodge, R., Brasington, J., & Richards, K. (2009). *In situ* characterization of grain-scale fluvial morphology using Terrestrial Laser Scanning. Earth Surface Processes and Landforms, 34(7), 954–968. 10.1002/esp.1780

14. Lane, S. N., Gentile, A., & Goldenschue, L. (2020). Combining UAV-Based SfM-MVS Photogrammetry with Conventional Monitoring to Set Environmental Flows: Modifying Dam Flushing Flows to Improve Alpine Stream Habitat. Remote Sensing, 12(23), 3868. 10.3390/rs12233868

15. Leduc, P., Peirce, S., & Ashmore, P. (2019). Short communication: Challenges and applications of structure-from-motion photogrammetry in a physical model of a braided river. Earth Surface Dynamics, 7(1), 97–106. 10.5194/esurf-7-97-2019

16. Legleiter, C. J. (2012). Remote measurement of river morphology via fusion of LiDAR topography and spectrally based bathymetry. Earth Surface Processes and Landforms, 37(5), 499–518. 10.1002/esp.2262

17. Marteau, B., Vericat, D., Gibbins, C., Batalla, R. J., & Green, D. R. (2017). Application of Structure-from-Motion photogrammetry to river restoration. Earth Surface Processes and Landforms, 42(3), 503–515. 10.1002/esp.4086

18. Mažeika, S., Sullivan, P., Watzin, M. C., & Hession, W. C. (2004). Understanding Stream Geomorphic State in Relation to Ecological Integrity: Evidence Using Habitat Assessments and Macroinvertebrates. Environmental Management, 34(5), 669–683. 10.1007/s00267-004-4032-8

19. Nemes-Kókai, Z., Borics, G., Csépes, E., Lukács, Á., Török, P., T-Krasznai, E., Bácsi, I., & B-Béres, V. (2024). Role of microhabitats in shaping diversity of periphytic diatom assemblages. Hydrobiologia, 851(4), 959–972. 10.1007/s10750-023-05336-x

20. Patil, S. B. (2023). Evaluation of Physico-chemical parameters in relation to fish production of Bhima River, M.S., India. *Ecology*, Environment and Conservation, 29(03), 1421–1424. 10.53550/EEC.2023.v29i03.063

21. Rabeni, C. F. (2000). Evaluating physical habitat integrity in relation to the biological potential of streams. Hydrobiologia, 422/423, 245–256. 10.1023/A:1017022300825

22. Su, H., Feng, Y., Chen, J., Chen, J., Ma, S., Fang, J., & Xie, P. (2021). Determinants of trophic cascade strength in freshwater ecosystems: A global analysis. Ecology, 102(7), e03370. 10.1002/ecy.3370

23. Taira, D., Mark, R. Y. Y., Hsiung, A. R., Jaafar, Z., & Todd, P. A. (2024). Fish responses to manipulated microhabitat complexity in urbanised shorelines. Journal of Applied Ecology, 61(5), 1095–1108. 10.1111/1365-2664.14644

24. Thoms, M., Scown, M., & Flotemersch, J. (2018). Characterization of River Networks: A GIS Approach and Its Applications. JAWRA Journal of the American Water Resources Association, 54(4), 899–913. 10.1111/1752-1688.12649

25. Wen, J., Chen, Y., Liu, Z., & Li, M. (2022). Numerical Study on the Shear Stress Characteristics of Open-Channel Flow over Rough Beds. Water, 14(11), 1752. 10.3390/w14111752

26. Westoby, M. J., Brasington, J., Glasser, N. F., Hambrey, M. J., & Reynolds, J. M. (2012). ‘Structure-from-Motion’ photogrammetry: A low-cost, effective tool for geoscience applications. Geomorphology, 179, 300–314. 10.1016/j.geomorph.2012.08.021

27. Zhao, C. S., Yang, Y., Yang, S. T., Xiang, H., Wang, F., Chen, X., Zhang, H. M., & Yu, Q. (2019). Impact of spatial variations in water quality and hydrological factors on the food-web structure in urban aquatic environments. Water Research, 153, 121–133. 10.1016/j.watres.2019.01.015

28. Zhou, N., Sheng, S., He, L.-Y., Tian, B.-R., Chen, H., & Xu, C.-Y. (2023). An Integrated Approach for Analyzing the Morphological Evolution of the Lower Reaches of the Minjiang River Based on Long-Term Remote Sensing Data. Remote Sensing, 15(12), 3093. 10.3390/rs15123093

29. Zhu, P., Pan, B., Li, Z., He, H., Hou, Y., & Zhao, G. (2023). Responses of biodiversity to microhabitat heterogeneity in debris flow gullies: Assessing the impact of hydrological disturbance. Science of The Total Environment, 902, 166509. 10.1016/j.scitotenv.2023.166509

